# A longitudinal naturalistic fMRI study of trait anxiety and cross-phase consistency in subjective fear signature expression

**DOI:** 10.64898/2026.07.24.740524

**Authors:** Chung-Lien W. Chen, Feng-Chun B. Chou, Po-Yuan A. Hsiao, Pin-Hao A. Chen

**Affiliations:** Department of Psychology, National Taiwan University, No. 1, Section 4, Roosevelt Road, Taipei 10617, Taiwan, ROC; Department of Psychology, The University of Texas at Dallas, 800 W. Campbell Road, GR41, Richardson, TX 75080–3021, USA; Graduate Institute of Brain and Mind Sciences, National Taiwan University, No. 1, Section 1, Jen Ai Road, Taipei 100, Taiwan, ROC; Neurobiology and Cognitive Science Center, National Taiwan University, No. 1, Section 4, Roosevelt Road, Taipei 10617, Taiwan, ROC

**Keywords:** trait anxiety, subjective fear, naturalistic viewing fMRI, predictive brain model, individual differences, cross-phase consistency

## Abstract

Persistent fear is a central feature of anxiety, but it remains unclear whether this persistence is more closely related to the consistency of conditioned-threat processing or to the consistency of subjectively experienced fear. Using a longitudinal naturalistic viewing fMRI paradigm, we examined whether trait anxiety was associated with cross-phase consistency in the expression of two validated fear-related neural signatures. Fortytwo participants completed two fMRI phases separated by approximately three months, viewing the same 18 naturalistic videos followed by a restingstate segment, resulting in 19 analysis segments for cross-phase analyses. We applied two pretrained signatures of subjective fear and threat conditioning, independently developed by separate research groups, to estimate their expression over time and examined how consistently each signature was reexpressed across phases within participants. We found that higher trait anxiety was significantly associated with greater cross-phase consistency in subjective-fear signature expression, with a significantly stronger association than for threat-conditioning signature expression. The anxiety–consistency association was also stronger for the subjective-fear signature than for a general negative-affect signature. This association became more prominent over longer temporal windows and was not detected in the average time series of the amygdala, ventromedial prefrontal cortex, insula, or hippocampus. These findings suggest that individual differences in trait anxiety are associated with the consistent cross-phase reexpression of distributed neural patterns related to subjective fear. Taken together, this study demonstrates how predictive brain models combined with longitudinal naturalistic viewing can provide a new approach for examining individual differences in trait anxiety across repeated experiences of the same naturalistic contexts.

## Introduction

Adaptive behavior depends on the ability to experience fear, which allows individuals to detect potential danger and organize appropriate responses (Adolphs, 2013; Mobbs et al., 2015; Steimer, 2002). However, effective adaptation also requires fear responses to be flexibly updated when the meaning of a stimulus or context changes (Bouton, 2004; Milad and Quirk, 2012; Vervliet et al., 2013). This flexibility is supported by neural systems that regulate and update learned fear responses during fear extinction, particularly through interactions between the medial prefrontal cortex, especially the ventromedial prefrontal cortex, and the amygdala (Milad and Quirk, 2002; Quirk, 2002; Quirk and Beer, 2006). Disruption of this regulatory balance may allow learned fears to persist, contributing to heightened anxiety and maladaptive responses to threat (Hariri et al., 2003; Kim et al., 2011; Marin et al., 2020). However, fear is not a unitary construct. LeDoux and Pine’s (LeDoux and Pine, 2016) two-system framework distinguishes explicit, subjective fear from implicit, defensive fear, separating subjectively experienced fear from automatic threat-related responses. Consistent with this distinction, meta-analytic evidence suggests that explicit and implicit fear rely on partly distinct neural pathways (Tao et al., 2021). Thus, anxiety may be sustained not only by exaggerated defensive reactivity, but also by the persistence of subjective fear, making subjectively experienced fear a key dimension for understanding anxiety.

Despite its importance for understanding anxiety, subjective fear has often been assessed using self-report questionnaires that ask individuals to evaluate their fear and avoidance in predefined situations (Baker et al., 2002; Caballo et al., 2015; Connor et al., 2001). Classic measurements such as the Fear Survey Schedule (FSS) ask individuals to rate their discomfort or anxiety toward a fixed set of stimuli, while later adaptations, such as the FSS-II for Older Adults (FSS-II-OA), incorporate fears particularly relevant to older adults and assess both fear intensity and interference with daily functioning (Geer, 1965; Kogan and Edelstein, 2004). These questionnaires have been valuable for characterizing phobic fears and examining how responses to particular stimuli relate to anxiety. However, because these questionnaires conceptualize fear primarily as a response to predefined stimuli, they are less suited to determining whether trait anxiety is associated with a broader property of subjective fear that extends across situations and over time.

Moving beyond predefined stimulus-based measures is particularly important because subjective fear in daily life is constructed across changing contexts rather than elicited by a single isolated trigger. As events unfold, fear may emerge from the integration of multiple cues, contextual information, memories, bodily states, and appraisals (Barrett, 2017; Critchley and Garfinkel, 2017; Meyer et al., 2019; Raber et al., 2019). Accordingly, individuals higher in anxiety may differ not only in the intensity of fear elicited by particular objects, but also in the consistency with which fear-related experiences are expressed across situations and over time (Beckers et al., 2023; Kuppens et al., 2010; Pawluk et al., 2021). Testing this possibility requires moving beyond fixed mappings between stimuli and responses and developing approaches that can capture subjective fear as it unfolds within richer and more dynamic contexts (Jääskeläinen et al., 2021; Saarimäki, 2021; Vanderwal et al., 2017). Naturalistic viewing paradigms, such as movie watching, provide an innovative approach to capture this complexity by exposing participants to dynamic streams of social, perceptual, and emotional information while preserving a relatively controlled stimulus sequence across individuals (Jääskeläinen et al., 2021; Vanderwal et al., 2017; Winkler and Appel, 2024). However, measuring subjective fear in these contexts remains challenging. Self-reports provide a direct assessment of subjective feelings, but they typically translate dynamic emotional states into predefined categories or scalar ratings and may disrupt the ongoing emotional experience (Hutcherson et al., 2005; Taschereau-Dumouchel et al., 2020; Winkler and Appel, 2024). If subjective fear is represented as a distributed and high-dimensional internal state, examining its consistency across situations and over time requires methods that can capture distributed patterns of neural activity in the brain.

Multivariate fMRI analysis provides such an approach by characterizing subjective fear from distributed whole-brain activity rather than from responses within any single region (Haxby et al., 2014; Horikawa et al., 2020; Taschereau-Dumouchel et al., 2020). Although early work on subjective fear emphasized the amygdala (Adolphs et al., 2005; Feinstein et al., 2011), subsequent fMRI studies have found that brain regions beyond the amygdala also contribute to subjectively experienced fear (Alpers et al., 2009; Hudson et al., 2020; Mobbs et al., 2007; Stark et al., 2007). More broadly, multivariate brain models have been used to identify sensitive and specific neural patterns underlying diverse psychological processes, including pain (Krishnan et al., 2016; Wager et al., 2013), reward (L. J. Chang et al., 2022), guilt (Yu et al., 2020), trust (Chen et al., 2021), and negative emotion (Chang et al., 2015). Extending this approach to fear, wholebrain pattern analyses have produced predictive models that generalize across individuals, including the threat-predictive signature (TPS), a multivariate brain model trained to distinguish conditioned threat from safety cues (Reddan et al., 2018) and the Visually Induced Fear Signature (VIFS), a multivariate brain model developed to predict the intensity of subjective fear (Zhou et al., 2021). Together, these two brain models provide complementary neural signatures indexing threat conditioning and subjective fear, allowing these two dimensions of fear to be examined separately in relation to anxiety. Despite their shared relevance to fear and anxiety, the two signatures show little spatial overlap. In the original VIFS validation study, their whole-brain pattern-weight maps showed negligible spatial similarity (*r* = .02, permutation *p* = .125), and cross-prediction between conditioned threat and subjective fear was limited (Zhou et al., 2021). Specifically, the VIFS did not reliably distinguish conditioned-threat (CS+) from safety (CS-) cues, whereas the threat-conditioning signature did not significantly predict subjective-fear ratings. Together, these findings provide empirical evidence that the two signatures differ in both their brain patterns and predictive profiles.

A key advantage of these multivariate models is that they can be applied beyond the original datasets or contexts in which they were developed (Woo et al., 2017). Recent work has shown that affective signatures trained using standardized stimuli can generalize to dynamic naturalistic experiences. For example, Chan et al. (Chan et al., 2020) trained neural classifiers of valence and arousal using affective pictures and successfully decoded continuous affective states during movie watching, demonstrating the feasibility of cross-context affective decoding. This approach is particularly useful for studying fear because it enables moment-to-moment estimation of affective states without interrupting the ongoing experience, unlike continuous self-report, which may alter emotional processing through introspection or task demands (Jolly et al., 2022; Meer et al., 2020; Vanderwal et al., 2017). Applying validated fear-related neural signatures to naturalistic viewing therefore creates an opportunity to revisit a central question in anxiety research: whether persistent fear reflects consistent fear-related neural pattern expression as emotional experience unfolds in complex contexts (Grupe and Nitschke, 2013; Robinson, 2019; Shackman and Fox, 2016). Although this question has largely been examined using conditioning and extinction paradigms, these designs provide limited insight into how fear-related states persist during more naturalistic experiences (Duits et al., 2015; Dunsmoor and Paz, 2015; Milad and Quirk, 2012).

Addressing this gap therefore requires tracking fear-related neural signatures over time while distinguishing threat-conditioning signature expression from subjective-fear signature expression, as anxiety-related persistence may involve either or both systems (Fanselow and Pennington, 2018; LeDoux and Pine, 2016; Taschereau-Dumouchel et al., 2020). Motivated by the proposed importance of flexible adaptation of fear responses for anxiety, we examined whether trait anxiety was associated with cross-phase consistency in subjective-fear signature expression when participants viewed the same videos again. We further examined whether a similar association was observed for the threat-conditioning signature, which was trained to distinguish conditioned threat from safety cues (Reddan et al., 2018). Events in naturalistic videos may cue threat meanings acquired through prior direct, observational, or informational learning, even in the absence of an experimentally imposed CS–US contingency (Olsson et al., 2007; Raber et al., 2019; Rachman, 1977). However, because the present paradigm did not experimentally manipulate threat learning, the threat-conditioning signature provides a complementary comparison rather than a direct test of anxiety-related differences in conditioned-threat learning.

Determining whether anxiety is associated with consistent neural signature expressions related to either subjective fear or conditioned threat also requires specifying when and where such consistency is expressed. Temporally, fear-related consistency may extend across an entire emotional episode, emerge within shorter local intervals, or operate at both time scales (Hudson et al., 2020; Kuppens et al., 2010; Zaki, 2013). At the brain regional level, an effect detected using distributed whole-brain models should also be compared with responses in canonical fear-related regions, including the amygdala, vmPFC, and insula (Fullana et al., 2016; Kragel et al., 2018; Phelps and LeDoux, 2005). In addition, because repeated viewing may involve memory and familiarity processes, we included the hippocampus as a complementary ROI to examine whether cross-phase consistency was primarily reflected in a region strongly implicated in episodic memory. Together, these considerations motivate a design that combines repeated naturalistic stimulation, validated fear-related brain models, temporal-scale analyses, comparison with canonical fear-related regions, and an additional hippocampal ROI analysis related to memory and familiarity (Chen et al., 2017; Nastase et al., 2019; Vanderwal et al., 2017).

To address these questions, the present study combined multivariate brain decoding with continuous movie viewing to examine how trait anxiety relates to cross-phase consistency of two distinct fear-related neural signatures: a subjective-fear signature and a threat-conditioning signature, each indexed by a validated multivariate brain model. Participants viewed the same set of naturalistic video clips during fMRI scanning at two time points separated by three months, allowing us to assess whether these neural signatures were consistently reexpressed when individuals encountered the same complex emotional contexts over time. We applied the subjective-fear and threat-conditioning models to derive moment-to-moment estimates of signature expression during each phase and quantified cross-phase consistency as the similarity between each participant’s model-expression time series across phases. We hypothesized that individuals with higher trait anxiety would exhibit greater cross-phase consistency in subjective-fear signature expression, reflecting more consistent reexpression of neural patterns related to subjective fear when encountering the same emotional contexts across the two phases. By contrast, we expected the association between trait anxiety and cross-phase consistency in threat-conditioning signature expression to be weaker. To characterize the temporal scale of this effect, we examined whether greater subjective-fear consistency in individuals with higher trait anxiety was prominent across the full time series or within shorter sliding windows. We expected this association to become more evident over longer temporal windows, suggesting that fear-related consistency emerges across sustained emotional contexts rather than isolated moments. We also tested whether the effect could be detected in the average time series of the amygdala, vmPFC, and insula, thereby assessing whether it was preferentially captured by distributed whole-brain patterns. Since repeated viewing may additionally involve memory and familiarity processes, we also included the hippocampus as a complementary ROI related to memory and familiarity. Together, this experimental design allowed us to test whether trait anxiety is more strongly associated with cross-phase consistency in subjective-fear signature expression than in threat-conditioning signature expression, whether this association is preferentially captured by distributed whole-brain patterns, and whether it can be distinguished from consistency in average brain regional activity. By separating subjective-fear and threat-conditioning signature expression within the same repeated naturalistic viewing paradigm, the present study aimed to clarify which form of fear-related neural expression is more closely associated with trait anxiety.

## Methods

### Participants

Seventy participants were recruited from National Taiwan University and completed Phase 1 of the fMRI experiment. Of these, 42 participants (all Asians) returned to complete Phase 2. Among these participants, the interval between the Phase 1 and Phase 2 scans ranged from 71 to 140 days (*M* = 103.8, *SD* = 16.7 days). Their mean age was 22.45 years (SD = 3.10, range = 18–33 years). Based on self-reported gender, 23 participants self-identified as male and 19 self-identified as female. All subsequent analyses were conducted using data from these 42 participants. All participants were screened for standard MRI contraindications before scanning. None of the participants reported a history of psychiatric or neurological disorders. All participants received monetary compensation and provided informed consent in accordance with the guidelines of the Research Ethics Center at National Taiwan University. This study was approved by the Research Ethics Center at National Taiwan University under approval number 202206HM013.

### Experimental procedure

#### Behavioral measurements: State-Trait Anxiety Inventory (STAI)

All participants completed the State–Trait Anxiety Inventory (STAI; (Spielberger et al., 1970) before fMRI scanning. The STAI is a widely used measure of two dimensions of anxiety: state anxiety, which reflects transient feelings of anxiety at a particular moment, and trait anxiety, which reflects a relatively consistent tendency to experience anxiety across situations and over time. Since the present study focused on individuals’ relatively consistent tendency to experience anxiety rather than temporary fluctuations, the subsequent analyses were based on the Trait Anxiety subscale (STAI-T). For subsequent analyses, all 42 participants were ordered according to their STAI-T scores from lowest to highest and assigned corresponding trait-anxiety ranks. Participants with lower STAI-T scores therefore received lower ranks, whereas those with higher scores received higher ranks. These rank-transformed values were used as a continuous predictor to test whether cross-phase consistency increased as trait anxiety increased.

#### Longitudinal naturalistic viewing fMRI paradigm

After completing the STAI, participants underwent a longitudinal naturalistic viewing fMRI paradigm consisting of two scanning phases separated by approximately three months. Each phase included 19 analysis segments, consisting of 18 video-viewing segments followed by one resting-state segment. During Phase 1, participants passively viewed 18 video clips covering a range of naturalistic contexts, including sports, social relationships, politics, travel, comedy, and nature. The 18 movie segments ranged from 16 to 228 TRs in duration (32 to 456 seconds; TR = 2 seconds), and the post-viewing resting-state segment contained 150 TRs (300 seconds). Detailed descriptions, sources, and individual durations of the movie segments are provided in Table S1. During Phase 2, the 42 returning participants completed the same procedure and viewed the identical sequence of video clips under the same scanning protocol. The final segment consisted of a 5-minute resting-state scan, during which participants viewed a fixation cross on the screen. Cross-phase consistency was calculated separately for each of the 19 segments. By keeping the stimulus content, presentation order, and scanning protocol constant across phases, this longitudinal repeated-viewing design allowed us to examine within-subject cross-phase consistency in neural responses to the same complex stimuli, as well as during the post-viewing resting-state segment (Figure 1).

**Figure 1.**
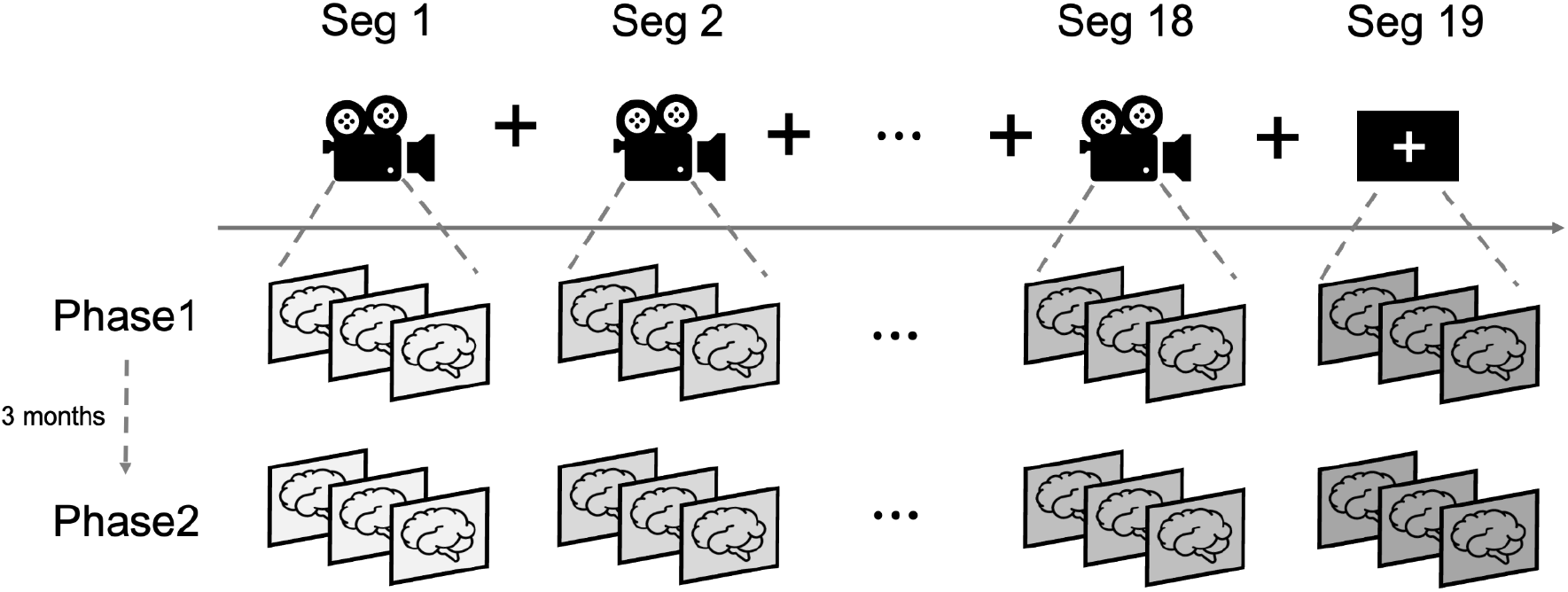
Longitudinal naturalistic viewing fMRI paradigm. Forty-two participants completed the same naturalistic viewing paradigm in two scanning phases separated by approximately three months. Each phase consisted of 19 analysis segments, including 18 video-viewing segments followed by one post-viewing resting-state segment. During Phase 1, participants passively viewed 18 videos presented in a fixed sequence while fMRI data were continuously acquired. During Phase 2, the same participants viewed the identical sequence using the same presentation timing and scanning protocol. Consecutive videos were separated by fixation intervals, and each phase concluded with a 5-minute resting-state scan.

### Image acquisition and preprocessing

MRI scanning was conducted using a 3 T Siemens Prisma scanner equipped with a 32-channel head coil at the NTU Imaging Center for Integrated Body, Mind and Culture Research. Structural images were acquired using a T1-weighted MPRAGE sequence (TR = 2300 ms, TE = 2.32 ms, voxel size = 0.9× 0.9 ×0.9 mm^3^, flip angle = 8^*°*^, FOV = 240 mm, 192 slices). Functional images were acquired using a T2*-weighted echo-planar imaging sequence (TR = 2000 ms, TE = 25 ms, voxel size = 3 ×3 ×3 mm^3^, flip angle = 75^*°*^, FOV = 240 mm, 40 slices). The preprocessing pipeline consisted of two stages. In the first stage, anatomical and functional images were preprocessed using fMRIPrep version 20.2.1 (Esteban et al., 2019). For the structural images, preprocessing included intensity nonuniformity correction and skull stripping using ANTs (Avants et al., 2009; Tustison et al., 2010). Brain tissue was then segmented into gray matter, white matter, and cerebrospinal fluid using FAST from FSL (Zhang et al., 2001). Finally, the T1-weighted images were nonlinearly normalized to the ICBM 152 template (MNI152NLin2009cAsym) at 2-mm resolution.

For functional images, preprocessing included motion correction, susceptibility distortion correction, and spatial normalization. A reference BOLD image was first generated and aligned to the T1-weighted reference using boundary-based registration (Greve and Fischl, 2009). Slice timing correction was applied, and BOLD time-series data were resampled to native and standard spaces with motion correction applied. Several confounding signals were also derived during preprocessing, including framewise displacement, DVARS, and global signals from cerebrospinal fluid, white matter, and whole-brain masks. To further characterize physiological noise, fMRIPrep generated component-based confound regressors using both temporal CompCor (tCompCor) and anatomical CompCor (aCompCor) (Behzadi et al., 2007). The tCompCor components were derived from the most variable voxels within a brain mask, whereas the aCompCor components were estimated from cerebrospinal fluid and white matter masks projected into native space. The functional data were subsequently spatially smoothed using a Gaussian kernel with a 6-mm full width at half maximum. The functional images used in the analyses were normalized to MNI152NLin2009cAsym space at 2-mm isotropic resolution using a single composite transform with Lanczos interpolation.

In the second stage, a voxelwise general linear model (GLM) was estimated separately for each participant.

The residualization model included an intercept, a linear temporal trend, 24 motion regressors (Friston, 2005), comprising the six realignment parameters, their squared values, their temporal derivatives, and the squared derivatives, as well as spike regressors identified using 3-*SD* cutoffs for the global mean signal and the mean frame-to-frame absolute difference. The CSF and white-matter mean signals and the aCompCor components generated by fMRIPrep were not included as regressors in this model. The residual time series from this model were retained for all subsequent analyses (Chang et al., 2018; L. Chang et al., 2022; Chen et al., 2020). At the participant level, all 42 participants who completed both scanning phases were retained in the longitudinal analyses. No additional participant-level exclusions were applied based on head motion or fMRI data quality, and no participant-level framewise displacement (FD) or DVARS threshold was used. Motion-related outlier volumes were addressed using the spike regressors described above.

### Cross-phase Consistency Analysis

#### Extraction of brain-model expression

In order to examine two distinct aspects of fear-related processing, we applied two validated multivariate brain models: a subjective-fear model and a threat-conditioning model. The subjective-fear model (Zhou et al., 2021), referred to as the Visually Induced Fear Signature (VIFS), was developed to predict the intensity of subjective fear from distributed whole-brain neural patterns. This model was trained using data from a discovery sample of 67 participants who viewed emotionally evocative images and rated the fear they experienced while imagining themselves in the situations presented, using a 5-point Likert scale. Using support vector regression, the VIFS reliably predicted self-reported fear in both cross-validation and independent validation samples. It also distinguished different levels of fear intensity with high accuracy, including 100% classification accuracy for differentiating high-fear ratings of 4–5 from low-fear ratings of 1–2. The model was characterized by a distributed pattern of cortical and subcortical contributions, including the amygdala, insula, periaqueductal gray, and medial prefrontal regions. Its performance across different participant samples and imaging systems further supported its generalizability as a predictive brain model of subjectively experienced fear.

The threat-conditioning model (Reddan et al., 2018), referred to as the threat-predictive signature (TPS) was developed to distinguish neural responses to conditioned threat and safety cues. This model was trained using whole-brain fMRI data from 68 participants who completed a threat-conditioning paradigm involving unreinforced CS+ and CS-stimuli. A linear support vector machine with leave-three-subjects-out crossvalidation achieved an average classification accuracy of approximately 77%. The model showed a sensitivity of 71%, a specificity of 84%, a positive predictive value of approximately 81%, and an area under the curve of 0.82. The threat-predictive pattern involved a distributed set of brain regions, including the prefrontal cortex, periaqueductal gray (PAG), insula, globus pallidus, and dorsal anterior cingulate cortex (dACC). These regions are commonly implicated in threat conditioning, supporting the use of this model as a measure of conditioned-threat processing.

We next applied both models to the preprocessed fMRI time series to estimate their moment-to-moment expression during naturalistic viewing. Both pretrained signature maps had 2-mm isotropic voxels but differed in image dimensions from the functional images. Before calculating pattern expression, the functional images and signature maps were resampled, when necessary, to a common 2-mm MNI152NLin2009cAsym brain mask. Voxel correspondence was determined from the source and reference image affines, with interpolation applied when resampling was required. Pattern expression was then calculated using the same masked voxel locations in the functional images and signature maps. For each fMRI volume, we computed the Pearson correlation between the participant’s wholebrain activation pattern and the unthresholded weight map of each model. This spatial correlation, referred to as pattern expression, indicates the degree to which the distributed neural pattern represented by a model is expressed in the observed brain activity (L. J. Chang et al., 2022). Higher positive values indicate stronger expression of the corresponding brain model, whereas lower or negative values indicate lower similarity to the model pattern. Applying this procedure across all volumes produced a continuous pattern-expression time series for both the subjective-fear and threat-conditioning models for each participant and scanning phase (Figure 2(A)).

**Figure 2.**
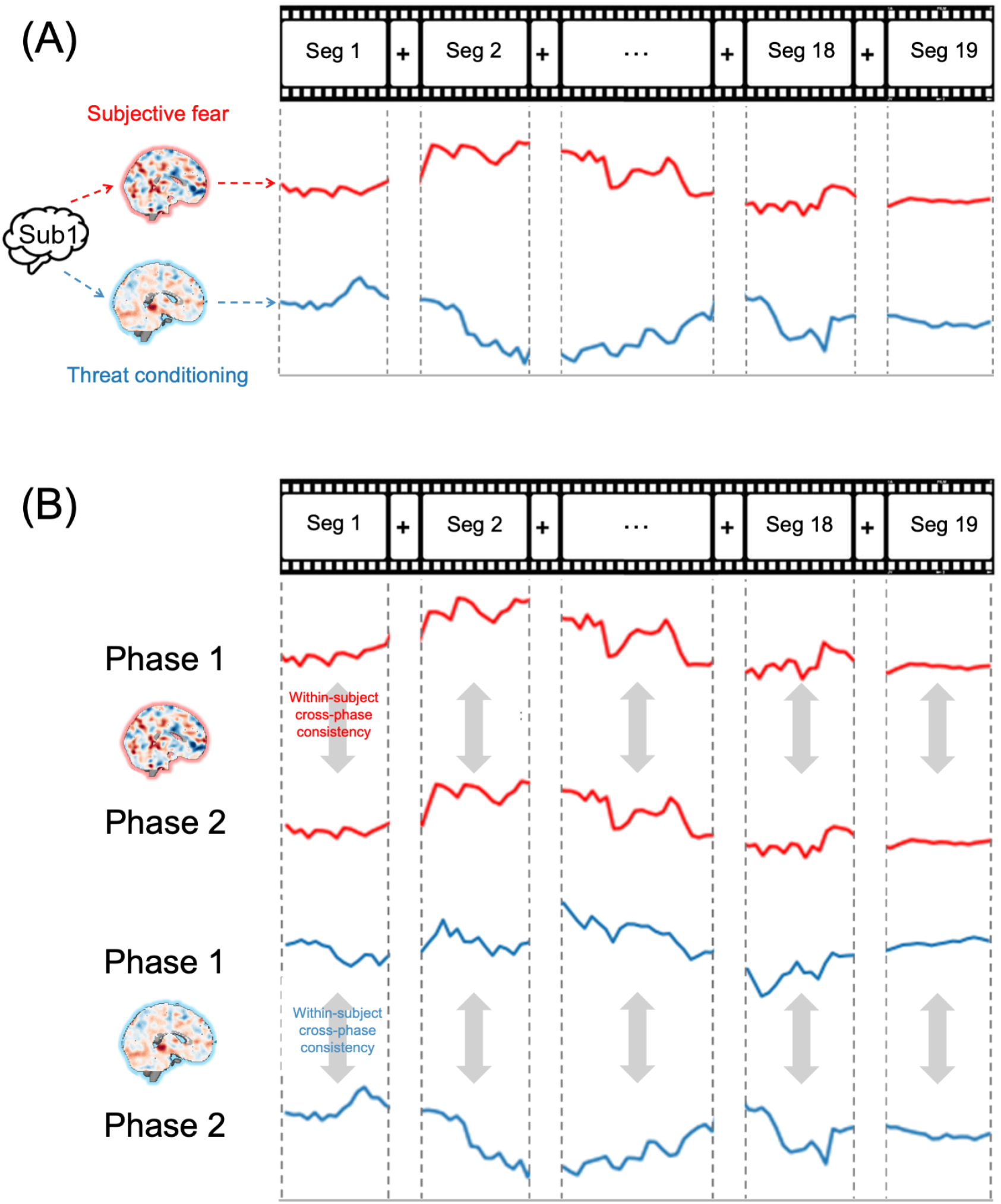
Extraction of brain-model expression and within-subject cross-phase consistency across nineteen analysis segments. (A) Each fMRI phase was divided into 19 analysis segments, including 18 video-viewing segments and one post-viewing resting-state segment. At each time point, the participant’s whole-brain activation pattern was spatially correlated with the unthresholded weight maps of the subjective-fear and threat-conditioning models. This procedure generated a continuous model-expression time series for each brain model within every segment. Solid red and blue lines illustrate the spatial Pearson correlation between the observed whole-brain activation pattern and each model weight map. (B) For each participant and analysis segment, model-expression time series were extracted separately from Phase 1 and Phase 2. Within-subject cross-phase consistency was computed as the Pearson correlation between the two expression time series from the corresponding segment, yielding one cross-phase consistency value for each participant, segment, and brain model.

#### Computing intrasubject cross-phase consistency value

To test whether higher trait anxiety was associated with greater cross-phase consistency in the expression of the two fear-related brain patterns, we examined how consistently each brain model was expressed within the same participant across Phase 1 and Phase 2. Each phase was divided into 19 analysis segments, including 18 video-viewing segments and one post-viewing resting-state segment. For each participant, model-expression time series were extracted separately for the subjective-fear and threat-conditioning models within each segment and phase. We then calculated the Pearson correlation between the Phase 1 and Phase 2 expression time series for the corresponding segment, yielding one cross-phase consistency value for each participant, segment, and model (Figure 2(B)). Higher cross-phase consistency values indicate that the decoded fear-related dynamics within a given segment were more similar across the two phases. Before subsequent analyses, cross-phase consistency values were rank-transformed separately within each of the 19 segments, and STAI-T scores were rank-transformed across participants.

We next examined whether trait anxiety was reliably associated with cross-phase consistency across the 19 analysis segments. Because cross-phase consistency was calculated separately for each segment, we used a two-stage grouped regression implemented with the Lm2 class in pymer4 (Jolly, 2018). In the first stage, a separate ordinary least-squares regression was fitted within each of the 19 analysis segments, with ranked consistency value as the dependent variable and ranked STAI-T score as the predictor. This analysis produced one anxiety–consistency slope for each segment. In the second stage, the segment-specific slopes were combined to estimate the overall association between trait anxiety and cross-phase consistency across the 19 segments. The resulting coefficient, *β*, was calculated as the unweighted arithmetic mean of the segment-specific slopes, with positive values indicating that higher trait anxiety was associated with greater cross-phase consistency. For *C* segments, *SE* was calculated as the sample standard deviation of the segment-specific slopes divided by 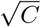, and *t* was calculated as *β*/*SE*, with *C* 1 degrees of freedom. First-stage variances were not used as inverse-variance weights. This approach allowed us to estimate the overall anxiety–consistency association across the full set of segments rather than relying on any single segment.

To provide an additional measure of effect magnitude, we also calculated a standardized coefficient for each segment. Ranked STAI-T scores and ranked cross-phase consistency values were standardized within each segment before fitting the same first-stage regression. Because this was a simple regression between two ranked variables, the resulting standardized coefficient was equivalent to Spearman’s correlation coefficient (*ρ*). These standardized coefficients provide a common scale for describing the direction and magnitude of the within-segment associations and were used to compare associations across the two brain signatures and four ROIs.

Finally, statistical significance was assessed using 1,000 nonparametric circular-shift permutations. In each permutation, the Phase 1 model-expression time series was held fixed. For each participant, analysis segment, and brain signature, a lag was sampled independently and uniformly from the integers 1 through *T* - 1, where *T* is the analyzed series length, and applied to that signature’s Phase 2 time series. Shifts were not matched across signatures; independently sampled lags could coincide by chance. Zero shifts were excluded, but small nonzero shifts were not otherwise excluded. For the sliding-window analyses described below, each shifted Phase 2 series was reused across all temporal window sizes within that permutation. Following each shift, cross-phase consistency was recalculated between the unchanged Phase 1 and shifted Phase 2 time series. Cross-phase correlations were Fisher- *z* transformed; for the sliding-window analyses, the Fisher-*z* values were averaged across overlapping windows. The resulting consistency estimates were then rank-transformed within each segment and submitted to the same two-stage grouped regression procedure used for the observed data. Using a random seed of 42, this process was repeated 1,000 times to generate separate null distributions of *β* coefficients for each brain signature and window size. Two-tailed permutation *p* values were calculated as *p* = *b*/*B*, where *b* was the number of permuted coefficients whose absolute values were greater than or equal to the absolute value of the observed coefficient and *B* = 1,000; no plus-one adjustment was applied. The Fear–Threat contrasts were evaluated using the same procedure with 1,000 permutations, with the mean within-segment slope difference used as the test statistic. Their null distributions were generated by independently shifting the two signatures and refitting the regressions. The full-series and sliding-window analyses used independently generated circular shifts.

#### Sliding-window consistency analysis

To determine whether the association between trait anxiety and cross-phase consistency in subjective-fear and threat-conditioning signature expression varied across temporal scales, we repeated the cross-phase consistency analysis using a sliding-window approach with progressively longer windows. For each participant, analysis segment, and brain model, the Phase 1 and Phase 2 model-expression time series were divided into windows with maximum window sizes ranging from 15 to 225 TRs in 15-TR increments (30 to 450 s). Within each window, cross-phase consistency was calculated as the Pearson correlation between the corresponding Phase 1 and Phase 2 expression time series. Windows were advanced along the time series in one-TR (2-s) increments. The resulting window-level correlations were Fisher-*z* transformed and then averaged within each participant and analysis segment to obtain a single cross-phase consistency estimate for each maximum window size.

The same 19 analysis segments were retained across all maximum window sizes. When the specified maximum exceeded the length of a segment, the full available time series was used for that segment. Because the shortest segment contained 16 TRs, 15 TRs was the only maximum in the original 15-TR to 225-TR analysis for which all segments contributed moving windows. As the maximum window size increased, more segments therefore contributed their full time series rather than moving windows. At 225 TRs, this applied to 18 of the 19 segments, with only Segment 13 (228 TRs) remaining long enough to contribute moving windows. The 225-TR estimate was therefore expected to approach the full-series estimate under this procedure (see Figure S1 in the Supplementary Materials for an illustration of the procedure at 15, 30, and 225 TRs).

For each maximum window size, consistency values were rank-transformed separately within each analysis segment, and ranked STAI-T scores were entered as the predictor. The association between trait anxiety and cross-phase consistency was estimated separately for the subjective-fear and threat-conditioning models using the same two-stage grouped regression implemented with the Lm2 class described above. The resulting *β* coefficient represented the average association between trait anxiety and cross-phase consistency across the 19 segments at each maximum window size. Statistical significance was assessed using the same nonparametric circular-shift permutation procedure. In each permutation, the Phase 2 model-expression time series was circularly shifted relative to the unchanged corresponding Phase 1 time series within each participant and analysis segment before window-level consistency was recalculated. This procedure disrupted the original cross-phase temporal alignment while preserving the temporal autocorrelation and distributional properties of each time series. The recalculated consistency values were rank-transformed and submitted to the same two-stage grouped regression. Two-sided *p* values were calculated by comparing the observed *β* coefficient with the permutation-derived null distribution. The result based on the full segment-level time series was presented alongside the sliding-window estimates as a reference for cross-phase consistency estimated using the complete available time series.

To address the increasing use of full-series substitution at longer maximum window sizes, we conducted an additional complete-window sensitivity analysis using maximum window sizes from 3 to 15 TRs (6 to 30 seconds; Figure S2). Within this range, all 19 segments contributed at least one complete window at every tested maximum, without shorter-than-maximum full-series substitution. The analysis otherwise followed the same procedure as the primary sliding-window analysis, including Fisher-*z* transformation and averaging of window-level correlations, within-segment rank transformation, the two-stage grouped regression, and 1,000 two-sided circular-shift permutations. Phase 1 remained fixed, whereas Phase 2 was shifted independently within each participant, segment, and signature. Within each permutation, the same shifted time series was used across all tested maximum window sizes before consistency values, ranks, and regression slopes were recalculated.

#### ROI-based analysis

To test whether the association between trait anxiety and cross-phase consistency in fear-related brain-model expression was specific to the distributed whole-brain pattern or could also be detected in the average time series of individual brain regions, we conducted a parallel ROI-based analysis. Specifically, we examined whether participants with higher trait anxiety also showed greater cross-phase consistency in regional fMRI time series, using the same participants, 19 analysis segments, and Phase 1–Phase 2 comparisons as in the whole-brain signature analyses described above.

ROI time series were extracted from a Neurosynth-based parcellation comprising 50 nonoverlapping brain regions (de la Vega et al., 2016). From this parcellation, we selected three a priori ROIs with strong theoretical relevance to fear and affective processing, including the amygdala, ventromedial prefrontal cortex (vmPFC), and insula. Because repeated viewing may also involve memory and familiarity processes, we additionally included the hippocampus as a complementary ROI associated with episodic memory. For each participant and ROI, regional time series were extracted separately for Phase 1 and Phase 2 within each of the 19 analysis segments. Cross-phase consistency was calculated as the Pearson correlation between the corresponding Phase 1 and Phase 2 regional time series, yielding one consistency value for each participant, segment, and ROI. These regional consistency values were then analyzed using the same rank transformation, two-stage grouped regression, and circular-shift permutation procedures described in Section 2.4.2.

#### Control analysis using a general negative affect brain signature

To examine whether the association between trait anxiety and cross-phase consistency in subjective-fear signature expression could reflect processes shared with general negative affect, we conducted an additional control analysis using the Picture-Induced Negative Emotion Signature (PINES; (Chang et al., 2015), a validated multivariate brain signature of general negative affect. PINES was independently developed to predict negative affective responses to emotionally evocative images and was not trained using the present dataset. Following the same procedures described above, PINES expression was calculated from the same preprocessed fMRI data for all 42 participants across the 19 segments. Using the same pattern-expression procedure described above, we computed the spatial Pearson correlation between the whole-brain activation pattern at each time point and the unthresholded PINES weight map, producing a continuous PINES-expression time series for each participant and phase. Cross-phase consistency was then calculated between corresponding Phase 1 and Phase 2 PINES-expression time series for each of the 19 analysis segments.

We applied the same within-segment rank transformation and two-stage grouped regression used in the primary analyses to estimate the association between trait anxiety and PINES cross-phase consistency. To compare this association with that observed for the subjective-fear signature, we calculated the within-segment difference between the subjective-fear and PINES anxiety–consistency slopes and averaged these differences across the 19 segments. Statistical significance was evaluated using 1,000 two-sided circular-shift permutations, with Phase 2 expression time series independently shifted relative to Phase 1 for each participant, segment, and signature. This procedure generated a null distribution for the subjective-fear– PINES slope contrast while disrupting the original cross-phase temporal alignment.

## Results

### Higher trait anxiety was associated with greater cross-phase consistency in subjective-fear signature expression

We first examined whether trait anxiety was associated with cross-phase consistency of subjective-fear and threat-conditioning signature expression across the full time series of the 19 analysis segments. The two-stage grouped regression showed a positive association for the subjective-fear signature, *β* = 0.097, *SE* = 0.040, *t*(18) = 2.409, *p* = 0.004. Participants with higher trait anxiety therefore showed greater cross-phase consistency of subjective-fear signature expression. The corresponding association for the threat-conditioning signature was not significant, *β* = -0.006, *SE* = 0.029, *t*(18) = -0.194, *p* = 0.864. A direct Fear–Threat comparison yielded a positive mean segment-level slope difference, *β* = 0.103, *SE* = 0.059, *t*(18) = 1.736, *p* = 0.042 (Figure 3A). All p-values were obtained from 1,000 two-sided circular-shift permutations. The interaction evaluates the observed slope contrast under disrupted cross-phase temporal alignment and does not by itself establish psychological specificity to fear.

**Figure 3.**
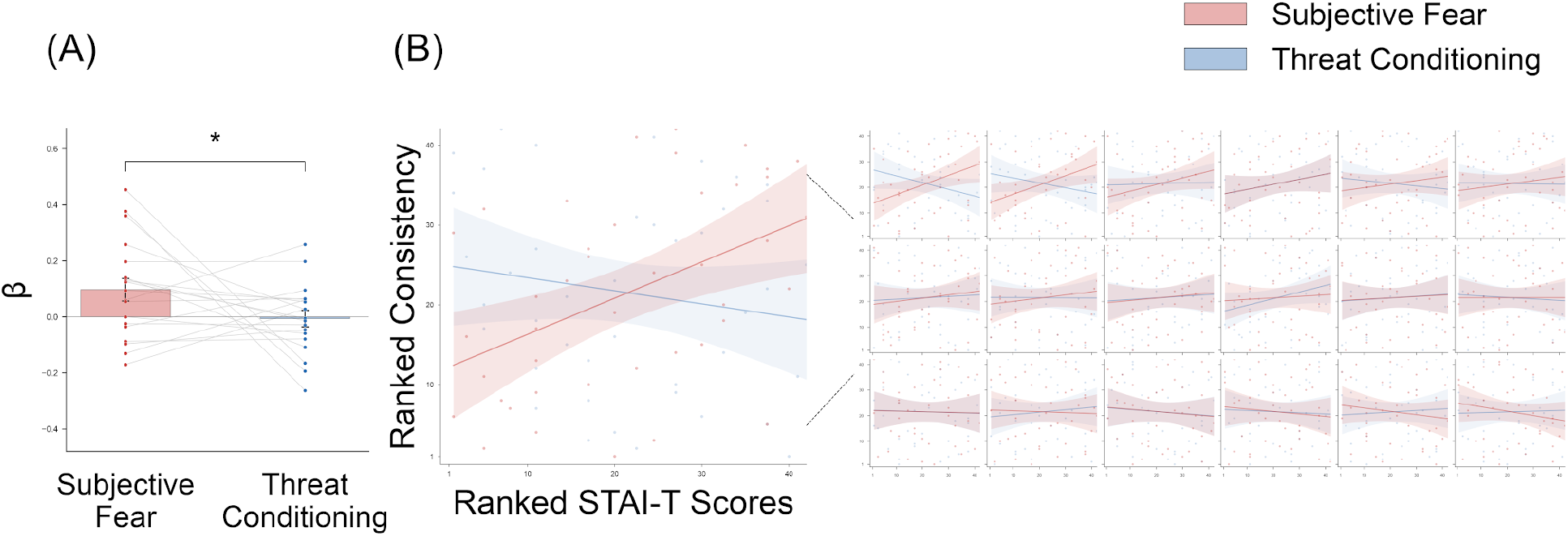
Trait anxiety is associated with cross-phase consistency of subjective-fear signature expression. (A) Associations between ranked STAI-T scores and cross-phase consistency are summarized across the 19 analysis segments, including 18 video-viewing segments and one post-viewing resting-state segment, for each brain signature. Bars represent the mean regression coefficient (*β*) across the 19 segments, dots represent segment-specific estimates, connected dots link estimates from the same analysis segment for the subjective-fear and threat-conditioning signatures, and error bars indicate *±*1 *SE*. The bracket denotes the direct Fear–Threat contrast, evaluated using 1,000 two-sided circular-shift permutations (*p* = .042; * *p* < .05). (B) For each analysis segment, ranked cross-phase consistency was plotted against ranked STAI-T scores separately for the subjective-fear and threat-conditioning signatures. Red represents subjective-fear signature expression, whereas blue represents threat-conditioning signature expression. Lines indicate fitted linear regressions with 95% confidence intervals, and points represent the 42 participants. Analysis segments are ordered according to the regression coefficient for subjective-fear signature expression.

We conducted several additional analyses to examine potential influences on the primary findings. First, we examined whether repeated viewing was associated with changes in mean signature expression across phases. Mean expression did not significantly differ between Phase 1 and Phase 2 for either the subjective-fear or threat-conditioning signature, and phase-to-phase changes in mean expression were not significantly associated with trait anxiety (Figure S3). Thus, we found no evidence of an overall phase-level change in signature expression or an anxiety-related change in mean expression. Because the post-viewing restingstate segment did not involve naturalistic video viewing, we examined whether the primary subjective-fear association remained when this segment was excluded. After restricting the analysis to the 18 movie segments, the association between trait anxiety and cross-phase consistency in subjective-fear signature expression remained positive and significant, *β* = .082, *SE* = .039, *t*(17) = 2.074, *p* = .025 (Figure S4). Finally, we examined whether the primary findings were influenced by any single participant or analysis segment using leaveone-out sensitivity analyses. The subjective-fear association remained positive after each of the 42 participant omissions (*β* = .077–.130) and each of the 19 segment omissions (*β* = .077–.112; Figure S5). The difference between the subjective-fear and threat-conditioning estimates also remained positive across all omissions. Together, these additional analyses indicate that the primary subjective-fear association was observed when the analysis was restricted to naturalistic viewing and was not driven by any single participant or analysis segment, while providing no evidence of an overall phase-level change in signature expression associated with repeated viewing.

### Higher trait anxiety was associated with greater subjective-fear consistency over longer temporal windows

To determine whether the association between trait anxiety and cross-phase consistency varied with the requested maximum window, we repeated the analysis using progressively longer sliding windows. The subjective-fear association was positive but nonsignificant at 15 and 30 TRs and became significant at 45 TRs, *β* = 0.079, *SE* = 0.041, *t*(18) = 1.942, *p* = 0.022.

The association remained significant across the longer tested maxima, reaching *β* = 0.097, *SE* = 0.040, *t*(18) = 2.407, *p* = 0.004, at 225 TRs. The full-series estimate was similar, *β* = 0.097, *SE* = 0.040, *t*(18) = 2.409, *p* = 0.004. By contrast, the association for the threat-conditioning signature was not significant at any tested maximum, with a full-series estimate of *β* = -0.006, *SE* = 0.029, *t*(18) = -0.194, *p* = 0.864. All p-values were obtained from 1,000 two-sided circularshift permutations (Figure 4). These results describe the associations observed at each requested maximum and do not constitute direct tests of differences between window-specific effects.

**Figure 4.**
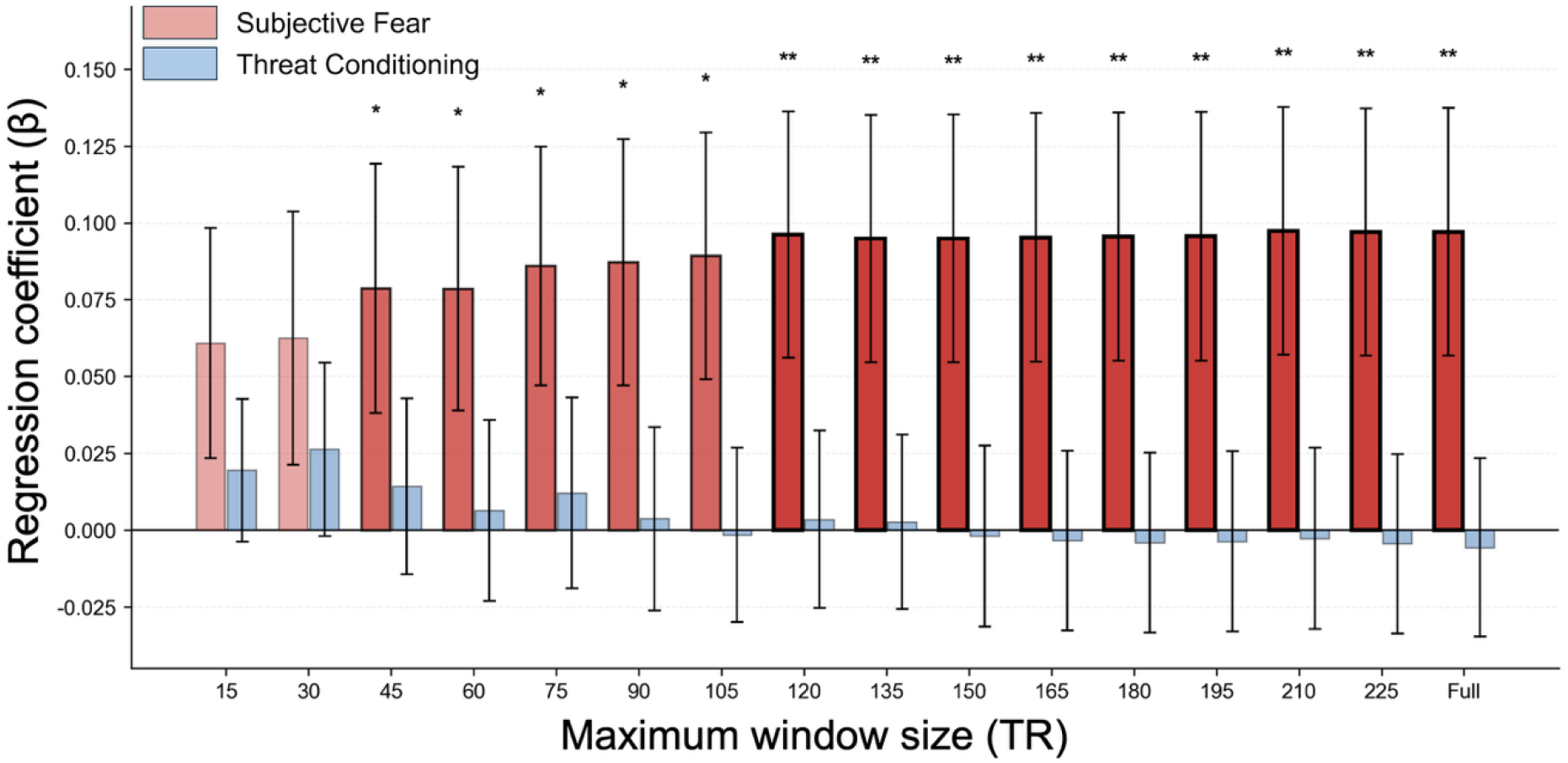
Trait anxiety was associated with greater cross-phase consistency in subjective-fear signature expression over longer temporal windows. Using the same analytic sample of 42 participants and 19 analysis segments, we examined whether the association between ranked trait anxiety and ranked cross-phase consistency varied across temporal window sizes. For each window ranging from 15 to 225 TRs, as well as for the full time series, associations were estimated separately for the subjective-fear and threat-conditioning models using the two-stage grouped regression with circular-shift permutation testing. Cross-phase consistency in subjective-fear signature expression showed a significant positive association with trait anxiety beginning at 45 TRs, and the association became more prominent over longer windows, approaching the full-series estimate. By contrast, cross-phase consistency in threat-conditioning signature expression was not significantly associated with trait anxiety at any window size. Bars represent regression coefficients, error bars indicate standard errors, and asterisks denote two-tailed permutation significance (\**p* < .05, \*\**p* < .01). Different color shades indicate levels of statistical significance.

The similarity between the 225-TR and full-series estimates should be interpreted in light of the maximum-window procedure. At 225 TRs, 18 of the 19 segments contributed their full available time series, whereas only Segment 13 contributed moving windows (Figure S1). The similarity between these estimates is therefore largely expected under the full-series substitution rule and does not provide independent evidence for a specific temporal-integration mechanism. We further conducted a complete-window sensitivity analysis using maximum window sizes from 3 to 15 TRs (6 to 30 s), for which all 19 segments contributed at least one complete window at every tested maximum without full-series substitution. No tested maximum showed a significant association with trait anxiety for either the subjective-fear or threat-conditioning signature (Figure S2). These results indicate that the complete-window analysis did not independently establish a significant anxiety–consistency association at these shorter temporal scales.

### The association between trait anxiety and subjective-fear consistency was detected in the distributed whole-brain pattern but not in selected ROIs

To determine whether the anxiety-related cross-phase consistency observed for subjective-fear signature expression could also be detected in individual brain regions, we conducted additional ROI-based analyses. We specifically examined the associations in the amygdala, vmPFC, and insula due to their roles in threat detection, fear learning, extinction, and emotional processing. The hippocampus was additionally included as a complementary ROI because repeated viewing may involve episodic memory or stimulus familiarity. The ROI and whole-brain signature analyses included the same 42 participants and 19 analysis segments. This yielded one cross-phase consistency value for each participant and segment for each ROI or brain signature. Cross-phase consistency and statistical significance were assessed using the same procedures applied in the whole-brain analyses, including the two-stage grouped regression and circular-shift permutation testing.

We found that trait anxiety was not significantly associated with cross-phase consistency in any of the selected ROIs. The associations were nonsignificant in the insula, *β* = .044, *SE* = .042, *t*(18) = 1.043, *p* = .215; vmPFC, *β* = .001, *SE* = .036, *t*(18) = .028, *p* = .976; amygdala, *β* = -.016, *SE* = .041, *t*(18) = -.393, *p* = .668; and hippocampus, *β* = -.033, *SE* = .036, *t*(18) = -.901, *p* = .355 (Figure 5). Together, these results indicate that the association observed for the distributed subjective-fear signature was not detected in the average regional time series of the selected ROIs. For comparison, the association was significant for the subjective-fear signature, *β* = .097, *SE* = .040, *t*(18) = 2.409, *p* = .004, but not for the threat-conditioning signature, *β* = -.006, *SE* = .029, *t*(18) = -.194, *p* = .864 (Figure 5). Within-segment standardized rankregression coefficients for all six measures are presented in Figure S6.

**Figure 5.**
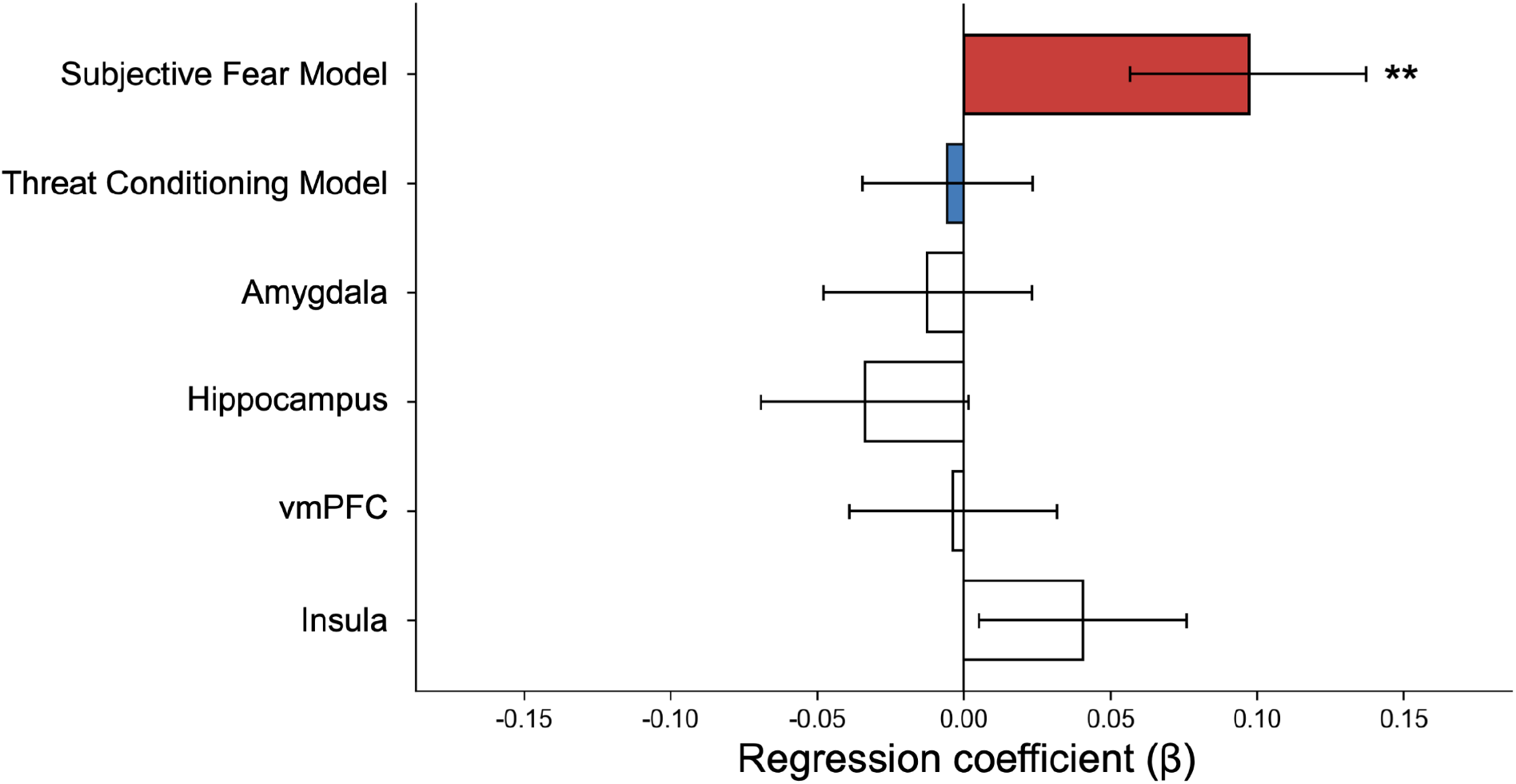
Comparison of associations between trait anxiety and cross-phase consistency across brain signatures and selected ROIs. Bars show the associations between trait anxiety and cross-phase consistency for the subjective-fear and threat-conditioning signatures and the four selected ROIs. A significant positive association was observed for the subjective-fear signature, whereas no significant associations were observed for the threat-conditioning signature or the insula, vmPFC, amygdala, and hippocampus. Error bars indicate *±*1 *SE* across the 19 segment-specific estimates. Red represents the subjective-fear signature, blue represents the threat-conditioning signature, and white represents the four individual ROIs. Statistical significance was evaluated using 1,000 two-sided circular-shift permutations. \*\**p* < .01.

### Subjective-fear signature consistency showed a stronger association with trait anxiety than negative-affect signature consistency

To examine whether the association between trait anxiety and cross-phase consistency in subjective-fear signature expression was similarly observed for general negative affect, we repeated the analysis using PINES (Chang et al., 2015), a validated multivariate signature of general negative affect. Across the 19 analysis segments, the mean anxiety–consistency slope was greater for the subjective-fear signature than for PINES (subjectivefear–PINES contrast: *β* = 0.144, *SE* = 0.048, *t*(18) = 3.026, *p* = .001; Figure S7). This comparison suggests that the association between trait anxiety and cross-phase consistency was stronger for the subjective-fear signature than for a general negative-affect signature. However, this comparison does not establish that the observed association is specific to fear or independent of broader negative affect.

## Discussion

In the present study, we examined whether trait anxiety was associated with cross-phase consistency in the expression of fear-related neural signatures during repeated naturalistic viewing. Participants viewed the same set of videos during two fMRI phases separated by approximately three months, allowing us to assess whether the two fear-related neural signatures were consistently reexpressed when individuals encountered the same emotional contexts again. These findings provide selective support for our primary hypothesis. Higher trait anxiety was associated with greater cross-phase consistency in subjective-fear signature expression, and this association was significantly stronger than for threat-conditioning signature expression. The temporal-scale analysis further supported our prediction that this association would become more prominent over longer windows. In addition, the effect was detected in the distributed whole-brain subjective-fear signature but was not detected in the average regional time series of the amygdala, vmPFC, insula, or hippocampus. Together, these findings suggest that trait anxiety is associated with the consistent reexpression of distributed neural patterns related to subjectively experienced fear, rather than with a general increase in the consistency of all fear-related neural processes.

The association between trait anxiety and subjective-fear signature expression offers a distinct perspective on the persistence of fear in individuals with high trait anxiety. Previous research has often emphasized impaired fear extinction as one mechanism underlying anxiety, whereby previously learned threat responses are not sufficiently reduced after repeated exposure (Davis et al., 2010; Fullana et al., 2016; Graham and Milad, 2011; Milad and Quirk, 2012). The present study did not directly measure extinction learning and therefore cannot determine whether the observed effect reflects an extinction deficit. However, our findings extend this broader perspective by suggesting that anxiety-related persistence may also involve the reliable reexpression of neural patterns related to subjective fear when individuals encounter the same emotional contexts again. Importantly, the effect reflected greater cross-phase consistency in subjective-fear signature expression rather than an overall increase in the level of expression. Trait anxiety may therefore be associated with greater consistency in subjective-fear-related neural expression across time. This interpretation is broadly in line with perspectives emphasizing sustained anticipatory processing, persistent negative appraisal, and difficulty disengaging from fear-related interpretations in anxiety (Davis et al., 2010; Grupe and Nitschke, 2013).

Trait anxiety was associated with greater cross-phase consistency in subjective-fear signature expression, and the direct fear–threat comparison indicated a greater anxiety–consistency association for the subjective-fear than for the threat-conditioning signature. This difference does not imply that threat conditioning is irrelevant to anxiety. The threat-conditioning signature was trained to distinguish conditioned threat from safety cues (Reddan et al., 2018), whereas the present naturalistic viewing paradigm did not experimentally manipulate threat acquisition, generalization, extinction, or updating, processes through which conditioning studies often examine anxiety-related differences (Beckers et al., 2023; Duits et al., 2015; Dunsmoor and Paz, 2015). Events in the videos could have cued threat meanings acquired through earlier direct, observational, or informational learning (Olsson et al., 2007; Raber et al., 2019; Rachman, 1977). However, we did not identify participant-specific threat cues or assess participants’ prior learning histories and intervening experiences, making it unclear whether the threat-conditioning signature reflected learned threat in these videos. The subjective-fear signature, by contrast, was developed to predict momentary subjective fear (Zhou et al., 2021). Its cross-phase consistency was associated with trait anxiety when participants viewed the same videos again, even in the absence of experimentally manipulated threat learning or extinction. These findings suggest that anxiety-related differences may become prominent in the repeated expression of subjective-fear-related neural patterns, not only in responses to changing threat contingencies. However, interpreting this association also requires considering whether the subjective-fear signature may capture processes shared with broader negative affect.

To examine this possibility, we repeated the analysis using PINES, a validated multivariate signature of general negative affect (Chang et al., 2015). The association between trait anxiety and cross-phase consistency was greater for the subjective-fear signature than for PINES across the 19 analysis segments, indicating that the observed anxiety–consistency relationship was not similarly expressed by a general negative-affect signature under the same analytical pipeline. However, this comparison does not establish fear specificity or statistically isolate subjective-fear signature expression from negative affect. Moreover, because the naturalistic clips were not independently validated as fear-inducing, other processes such as arousal may also contribute to the observed pattern. We therefore interpret our findings as an association between trait anxiety and cross-phase consistency in subjective-fear signature expression.

The temporal-scale analyses further showed that the association between trait anxiety and cross-phase consistency in subjective-fear signature expression became more prominent over longer temporal windows. The association was not significant at the shortest windows of 15 and 30 TRs but became significant at 45 TRs and remained significant across the longer tested maxima. One possible interpretation is that longer windows better capture emotional dynamics that unfold as perceptual, contextual, and narrative information accumulates over time. Continuous naturalistic perception depends on information integration across multiple temporal scales, with higher-order systems showing relatively long temporal integration windows (Chang et al., 2021; Hasson et al., 2008; Lerner et al., 2011; Soltani et al., 2021; Yeshurun et al., 2021). Shorter windows may be more sensitive to transient fluctuations in emotional processing, whereas longer windows may better capture sustained aspects of subjective emotional experience (Asutay et al., 2021; Kuppens and Verduyn, 2017). This interpretation is also broadly consistent with theoretical distinctions between sustained anxiety and brief phasic fear responses (Alvarez et al., 2011; Davis et al., 2010). However, this temporal pattern should be interpreted cautiously. The 225-TR estimate was largely based on full-series values because most segments were shorter than this window, and the complete short-window analyses did not show significant associations. Moreover, significant effects at longer but not shorter windows do not necessarily indicate a significant difference between temporal scales. Thus, the present findings suggest that the anxiety–consistency association was more apparent over longer time windows, but they do not establish a specific temporal-integration mechanism or timescale.

The ROI analyses provide a complementary perspective on the distributed nature of this association. Trait anxiety was not significantly associated with cross-phase consistency in the average time series of the amygdala, vmPFC, insula, or hippocampus. Prior work has implicated these regions in fear learning, extinction, emotional processing, and memory, although their involvement varies across paradigms and phases of learning (Fullana et al., 2018; Milad et al., 2007; Phelps et al., 2004). These earlier studies typically examined regional responses during experimentally defined fear acquisition, extinction, or recall, whereas the present study examined the consistency of regional dynamics when participants encountered the same naturalistic contexts again. The absence of significant ROI associations therefore does not imply that these regions are uninvolved in fear, anxiety, memory, or extinction. Nor do nonsignificant ROI associations by themselves establish a significant difference between regional and whole-brain measures. Instead, the ROI results provide a complementary reference for the association detected in the distributed subjective-fear signature. Subjective fear may involve the integration of perceptual information, bodily signals, contextual meaning, memory, and appraisal across distributed neural systems (Taschereau-Dumouchel et al., 2020; Zhou et al., 2021), consistent with the observation that the anxiety–consistency association was detected in distributed subjective-fear signature expression but not in the average regional dynamics of the selected ROIs.

More broadly, the repeated naturalistic viewing design combined with pretrained predictive brain models provides an alternative approach for examining the consistency of affect-related neural dynamics across time. Naturalistic paradigms preserve continuous and time-varying psychological experiences while eliciting corresponding neural dynamics across complex contexts (Chen et al., 2020; Finn et al., 2018; Hasson et al., 2004; Saarimäki, 2021; Vanderwal et al., 2019). By presenting the same stimulus sequence under the same scanning protocol across two phases, the present design allowed us to quantify within-person cross-phase consistency rather than treating changes across repeated viewing solely as noise. Predictive brain models further enabled us to estimate fear-related signature expression at each fMRI volume without repeatedly interrupting the ongoing experience with explicit ratings (Chen et al., 2020; Chen and Qu, 2021; Kim et al., 2022, 2024; Woo et al., 2017). Although selfreport remains essential for establishing subjective experience, continuous ratings and repeated probes can introduce attentional and cognitive demands that alter affective experiences during naturalistic viewing (Isik and Vessel, 2019; Jolly et al., 2022; Wagner et al., 2021). The present approach therefore extends the use of predictive brain models beyond estimating momentary psychological-state-related expression to examining how consistently model-derived neural dynamics are reexpressed within individuals across repeated experiences. The present study may provide a complementary approach for investigating stable individual differences in affective processing under naturalistic conditions.

Several limitations should be considered when interpreting these findings. First, applying pretrained signatures to naturalistic narratives necessarily involves a degree of construct uncertainty because complex videos simultaneously engage perceptual, cognitive, social, and affective processes (Saarimäki, 2021). Importantly, ecological richness and construct specificity represent separate considerations. Naturalistic viewing preserves the continuous and context-dependent structure of ongoing experience, whereas the expression of a pretrained neural signature remains a modelbased estimate of the construct it was developed to predict. As discussed above, the videos were not independently validated as fear-inducing, and the PINES comparison does not establish that the observed association is specific to fear. We also did not collect time-resolved reports of subjective fear during scanning. Consequently, moment-to-moment fluctuations in signature expression could not be directly compared with participants’ subjective experiences. The resulting expression values should therefore be interpreted as model-based estimates of neural patterns related to subjective fear and threat conditioning rather than as direct measurements of isolated mental states. Future studies could combine model-based decoding with retrospective continuous ratings or experience-sampling procedures designed for naturalistic paradigms to more directly establish how signature expression relates to ongoing subjective experience.

Second, repeated viewing may introduce familiarity, memory, or habituation effects that could influence neural responses across phases. Repeated exposure to the same stimuli can reduce emotional responses, and such habituation may vary across individuals. To examine this possibility, we compared the mean signature expression between the two phases. We found no significant phase difference for either the subjective-fear or threat-conditioning signature, and changes in signature expression were not significantly associated with trait anxiety. However, these null results do not establish the absence of habituation or other repeated-exposure effects. Our cross-phase consistency measure captures the similarity of signature-expression dynamics across phases rather than changes in their overall level. Repeated exposure could therefore alter the magnitude or subjective experience of emotional responses without necessarily reducing cross-phase consistency. In addition, the hippocampal analysis provided no evidence that trait anxiety was associated with hippocampal cross-phase consistency, although memory-related contributions distributed across other brain regions cannot be ruled out. We also examined whether the post-viewing resting-state segment influenced the main finding. After excluding this segment, the association between trait anxiety and subjective-fear signature consistency remained positive and significant across the 18 movie segments. Thus, the primary association was also observed when the analysis was restricted to naturalistic viewing.

Third, the final sample of 42 participants limits the precision and generalizability of our findings. In the leave-one-out analyses, the subjective-fear association remained positive after removing each participant or segment, and the difference between the subjective-fear and threat-conditioning estimates also remained positive. These results suggest that the main findings were not driven by a single participant or segment, although they should not be considered an independent replication. In addition, our sample consisted of nonclinical participants and therefore reflects continuous variation in trait anxiety rather than clinically diagnosed anxiety. Future studies with larger and clinical samples are needed to determine whether cross-phase consistency in subjective-fear signature expression generalizes across populations. Further work could also examine whether this neural consistency is related to clinically relevant outcomes, such as symptom persistence or treatment response.

In summary, using a longitudinal naturalistic viewing paradigm, we found that higher trait anxiety was associated with greater cross-phase consistency in subjective-fear signature expression. This association was observed for the subjective-fear signature but not for the threat-conditioning signature and became more prominent when consistency was estimated over longer temporal windows. The association was also detected in the distributed subjective-fear signature but not in the average time series of the selected fear-related and memory-related ROIs. Together, these findings suggest that cross-phase consistency in distributed subjective-fear-related neural expression may represent a complementary dimension for understanding individual differences in trait anxiety.

